# B-cell activating factor plays a critical role in CAR-T cell-associated cytokine release syndrome

**DOI:** 10.1101/2025.11.22.689920

**Authors:** Claire Fritz, Leland Metheny, David Wald, Paolo Caimi, Reshmi Parameswaran

## Abstract

Cytokine release syndrome (CRS) is the most common and potentially life-threatening toxicity associated with CAR-T cell therapy and is related to a heightened immune effector state. In this work, we identified a novel role of the pro-tumorigenic cytokine B-cell activating factor (BAFF) in its pathophysiology. First, we observed that patients who experienced CAR-T cell-related CRS have elevated serum BAFF levels that coincide with elevated IL-6. Mechanistically, we show that IFN-γ, produced by activated CAR-T cells, stimulates monocytes to release BAFF, which induces the expression of CRS-related cytokines from monocytes. Monocytes derived from CRS patients express BCMA, which is further induced by IFN-γ stimulation. Neutralization of BAFF with belimumab significantly reduces production of various CRS and ICANS-related cytokines without impairing CAR-T cell activation or killing. Overall, we demonstrate that BAFF plays a critical role in CAR-T-cell-related CRS, and its neutralization may be a novel strategy for treating both CRS and ICANS.

**Graphical abstract:** 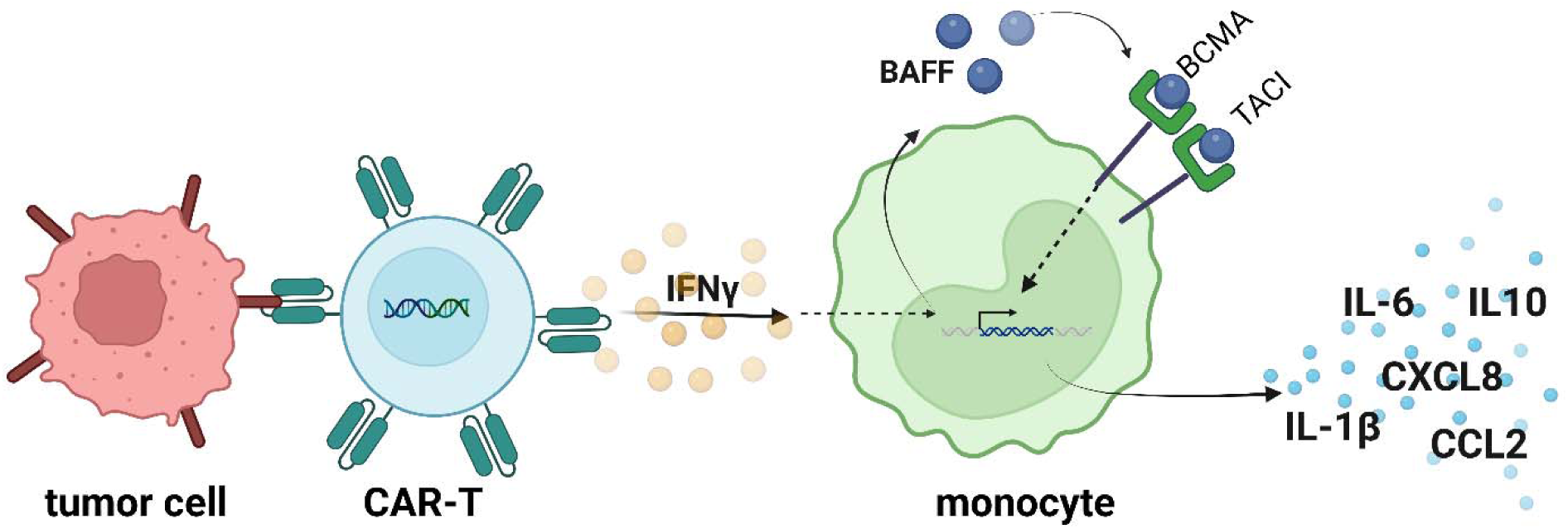

## Introduction

Despite the recent clinical success of chimeric antigen receptor (CAR)-T cell therapy in treating previously incurable relapsed and refractory hematological malignancies, toxicities remain a major concern.^1^ Cytokine release syndrome (CRS) and Immune Effector Cell-Associated Neurotoxicity Syndrome (ICANS) are the most common toxicities associated with CAR-T cell therapy. CRS is characterized by fever, hypotension, and hypoxia accompanied by supraphysiological levels of inflammatory cytokines.^2,3^ Severe cases present with capillary leak syndrome, multi-organ failure, and disseminated intravascular coagulation.^4^ Depending on the clinical trial, the incidence of CRS ranges from 57-93% with severe cases comprising 13-32% of cases and often requiring intensive care management.^5^ In addition to the risk to patient safety, the management of CRS increases healthcare resource utilization and the financial burden on patients.^6,7^ Considering the rapid expansion of CAR-T cell utilization in the clinic and high prevalence of these adverse events, recent studies are aimed at identifying predictive biomarkers, elucidating the mechanisms, and developing mitigation strategies for CAR-T cell-related CRS and ICANS.^3,8–16^

The underlying pathophysiology of CAR-T cell-associated CRS is generally attributed to the heightened immune effector state following CAR-T cell infusion. After infusion, CAR-T cells are trafficked to the site of tumor cells. CAR-T cells are activated and highly proliferative upon tumor antigen recognition. The heightened levels of the CAR-T-derived cytokines interferon-γ (IFN-γ), granulocyte-macrophage colony-stimulating factor (GM-CSF), and tumor necrosis factor α (TNFα), in addition to damage-associated molecular patterns (DAMPs) released from pyroptotic tumor cells, result in the recruitment of circulating monocytes, which have been recently identified as the central cellular mediator of the CRS cascade.^3,4,17^ Monocytes and activated macrophages are stimulated to produce supraphysiologic levels of pro-inflammatory cytokines and chemokines, including IL-6, IL-10, IL1β, CXCL8, and CCL2.^4,17,18^ The resulting pro-inflammatory feed-forward cycle and systemic inflammation leads to endothelial cell activation and vascular leakage and the symptoms of CRS. IL-6 is considered the most important driver of this process. As such, its neutralization with tocilizumab is standard of care, along with steroids and supportive care.^19^

To elaborate upon the pathophysiology of CAR-T-related CRS, we explored a previously unidentified role of the cytokine B-cell activating factor (BAFF) in this pathway. BAFF is a tumor necrosis factor (TNF) family cytokine critical for B-cell development and survival and is well known as a pro-tumorigenic factor that is elevated in the serum of patients with B-cell cancers.^20,21^ BAFF is produced by various cell types in the microenvironment, including monocytes, macrophages, stromal cells, and dendritic cells and binds to three receptors, the BAFF receptor (BAFF-R), transmembrane activator and CAML interactor (TACI), and B-cell maturation antigen (BCMA), to induce pro-survival and inflammatory signaling in healthy and malignant B-cells.^22,23^ Because of its prominent role in B-cell tumor microenvironments, we hypothesized that BAFF may contribute to the CRS cascade.

In this work, we identify a novel role of BAFF in the pathophysiology of CAR-T-cell-associated CRS. First, we establish the clinical relevance of BAFF in CRS by demonstrating that serum BAFF levels spike similarly to known cytokine mediators of CAR-T-associated CRS and are not elevated in CAR-T cell patients who did not experience CRS. To elucidate the role of BAFF in CRS pathophysiology, we first identified that monocytes are the primary producers of BAFF in this tri-culture and that stimulation of monocytes with IFN-γ results in increased BAFF production. Further, both TACI and BCMA are expressed on THPs, healthy monocytes, and monocytes derived from CAR-T-CRS patients, and stimulation with IFN-γ upregulates the expression of BCMA on these monocytes. Next, we found that BAFF induces the expression of various CRS-related cytokines in monocyte cells.

Neutralization of BAFF and BCMA in a tri-culture system of monocytes (THPs), cancer cells (Raji or Jeko), and CD19 CAR-T cells results in a substantial decrease in levels of the classical cytokines and chemokines involved in the CRS cascade, including IL-6, CXCL8, CCL2, GM-CSF, IL-1β, and IL-10. In a CAR-T CRS in vivo model, we found that serum BAFF levels significantly increased following CAR-T cell administration and that expression of key CRS cytokines showed a reducing trend with BAFF neutralization in vivo. Critically, BAFF neutralization with belimumab or atacicept does not impair CAR-T cell activation, degranulation, or cytotoxic activity.

## Results

### Patients who experience CAR-T-related CRS have elevated levels of serum BAFF

We analyzed serum cytokine levels of 11 CAR-T treated patients, seven of whom developed CRS and 4 who did not develop clinical CRS. We included serum levels before CAR-T infusion and at various timepoints thereafter, depending on sample availability. Our data shows that serum BAFF levels in CAR-T patients who experienced CRS showed a trend similar to IL-6 elevation (Figure 1). Although Patient 8 was not reported to experience clinical CRS, serum IL-6 levels peaked post-CAR-T infusion, so this patient was included in our analysis of CRS-positive patients. In patients who did not experience CRS after CAR-T infusion, serum BAFF levels did not show an appreciable increase (Figure 2). The serum BAFF levels were significantly higher than either IL-6 or IFN-γ at baseline, as B-cell cancer cells are known to secrete BAFF.^20^ However, we repeatedly saw the trend in these patients that serum BAFF levels increased even further following CAR-T cell administration. Further, the levels decreased toward baseline levels thereafter, similarly to the trend of IL-6 and IFN-γ (Figure 1). A proliferation-inducing ligand (APRIL), a related cytokine to BAFF, followed the same trend, though its serum levels were low compared to BAFF (Supplementary Figure 1). Furthermore, other CRS-related cytokines, CXCL8, CCL2, GM-CSF, IL-1β and IL-10 demonstrated a similar trend, peaking at a similar timepoint in these patients (Supplementary Figures 1,2).

**Figure 1:**
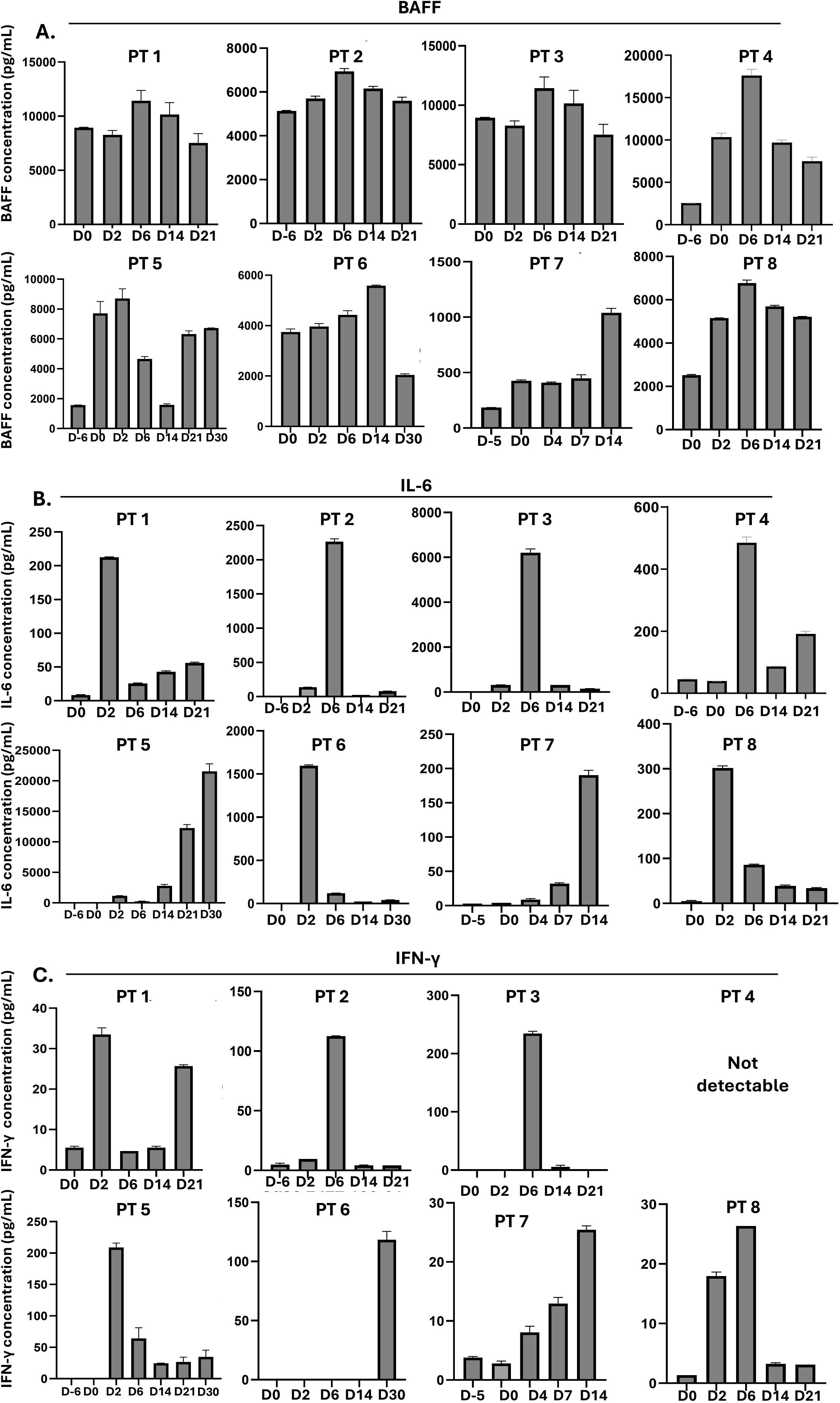
Serum BAFF is elevated in CAR-T patients who experience CRS. Serum from CRS+ patients at various timepoints after CAR-T-cell infusion (D=day post CAR-T-cell infusion) was subjected to Multiplex or Luminex cytokine analysis. ’PT’ is de-identified patient number.

**Figure 2:**
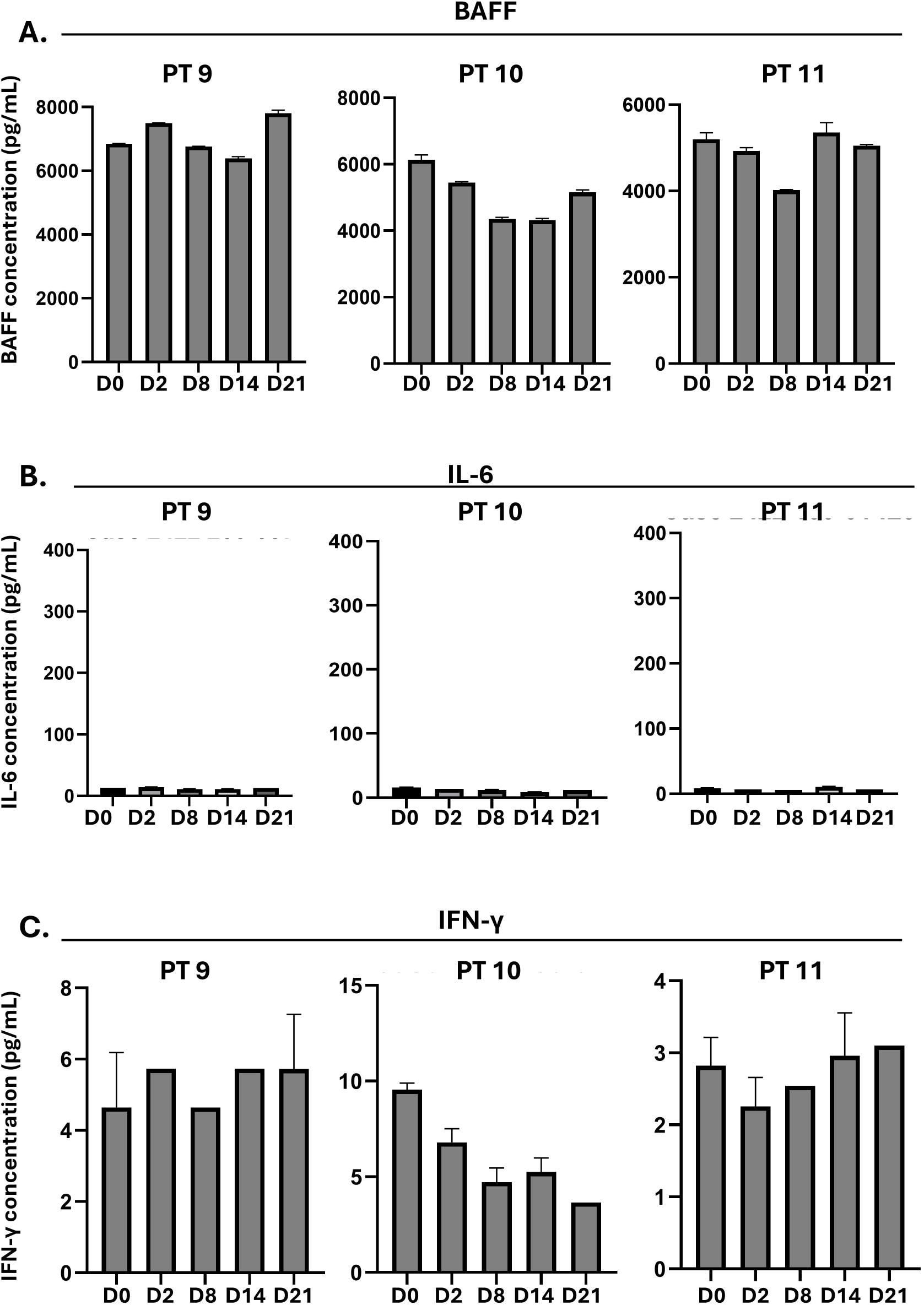
Serum BAFF does not increase in patients who did not experience CAR-T related CRS. Serum from patients who received CAR-T cell therapy and were not diagnosed with CRS at various timepoints after CAR-T-cell infusion (D=day post CAR-T-cell infusion) was subjected to Multiplex or Luminex cytokine analysis.

In the set of patients that we analyzed, 8 had diffuse large B-cell lymphoma (DLBCL) and were treated with commercial axicabtagene ciloleucel (Supplementary Figure 3). One patient with plasma cell leukemia was treated with LMY-029 (NCT05312801), and patients with follicular lymphoma, DLBCL, and mantle cell lymphoma were treated with UF-Kure19 (NCT05400109) (Supplementary Figure 3). The CRS grades of the patients included in our analysis ranged from 1-3: three patients had grade 1, two patients had grade 2, and 3 patients experienced grade 3 CRS (Supplementary Figure 3).

### BAFF is predominantly produced by monocytes

Upon discovering that serum BAFF levels increase in CAR-T-treated CRS patients, we were interested in identifying which cell types contribute to this increased BAFF and other CRS-related cytokines after CAR-T cell infusion. To this end, we cultured Jeko, THPs, and CAR-T cells alone and in combination with a 1:1 or 1:1:1 ratio. Each cell type produced BAFF independently, with THP cells secreting the highest amount at baseline. The combination of CAR-T cells and monocytes yielded the greatest increase in secreted BAFF (Figure 3). IL-6 was minimally produced by the combination of cancer and CAR-T cells. The co-culture of CAR-T cells with THPs significantly increased IL-6 and IL-1β secretion relative to any cell type alone or other combinations. Both cytokines increased even further with the addition of cancer cells to the co-culture. At a baseline, CAR-T cells were the only cell type that secreted a low level of IFN-γ or GM-CSF independently. The addition of either cancer cells or THPs to CAR-T cells substantially increased the production of both cytokines. Furthermore, APRIL was released at lower levels than BAFF, as we saw in our patient serum data. Similarly to BAFF, each cell type secreted APRIL alone, with THPs producing the most. The combination of all the cell types significantly increased APRIL production. CXCL8 and CCL2 were minimally produced by THP cells alone, but their secretion was significantly increased by the combination of CAR-T cells with THPs or all three cell types (Figure 3).

**Figure 3.**
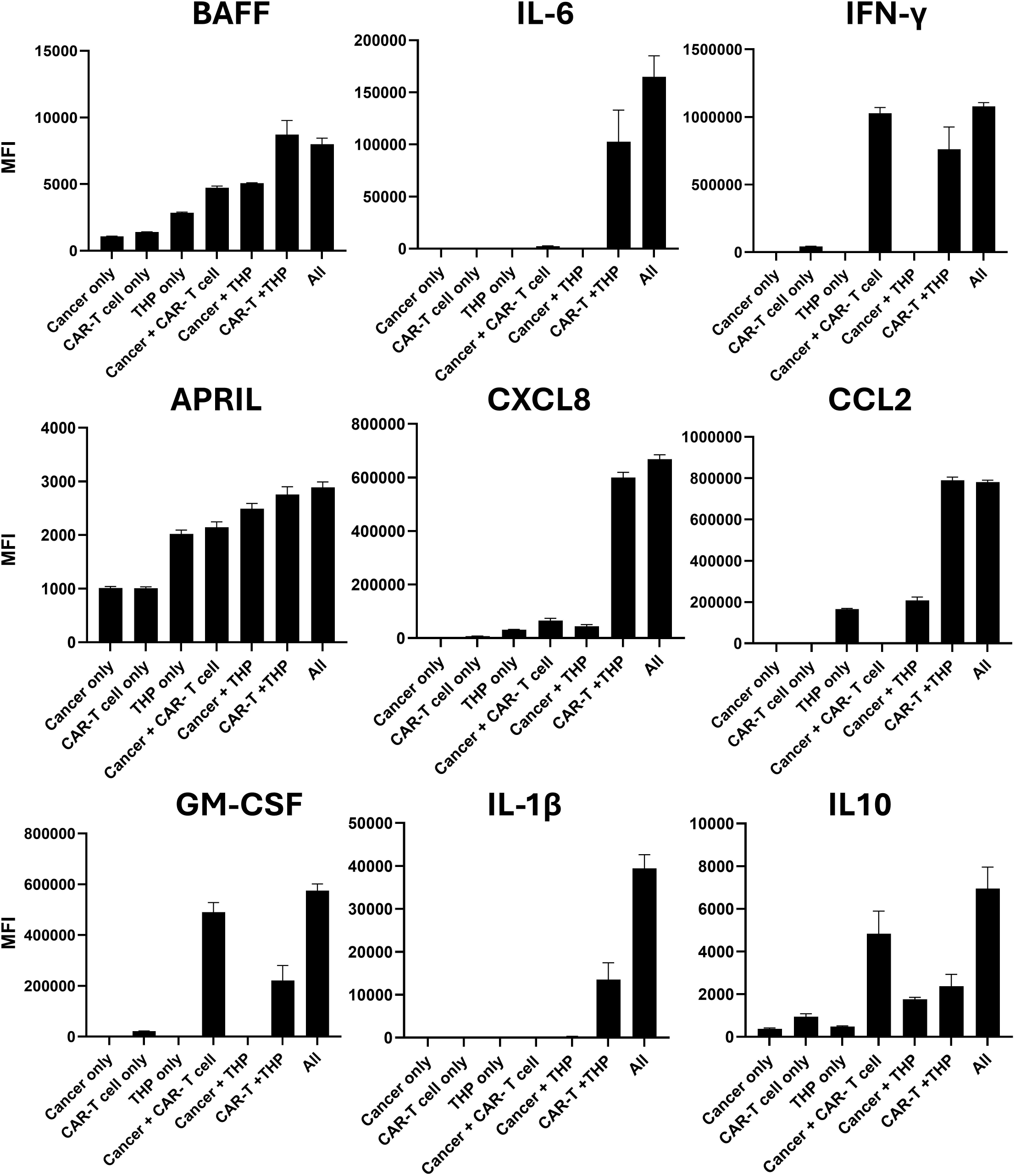
Contributions of cell types to cytokine release. Jeko-1 cells, CD19 CAR-T cells, and THP-1 cells were cultured alone or in every combination at a 1:1:1 ratio for 24h, when the supernatant was harvested for cytokine analysis by Multiplex flow cytometry. This data represents 2 independent experiments.

### IFN-**γ** induces BAFF secretion and BCMA expression on monocytes

To investigate a link between CAR-T activation and elevation of BAFF in the CRS setting, we stimulated monocytes with recombinant IFN-γ, a predominant cytokine produced post CAR-T activation, and saw an increase in BAFF secretion (Figure 4A).

**Figure 4.**
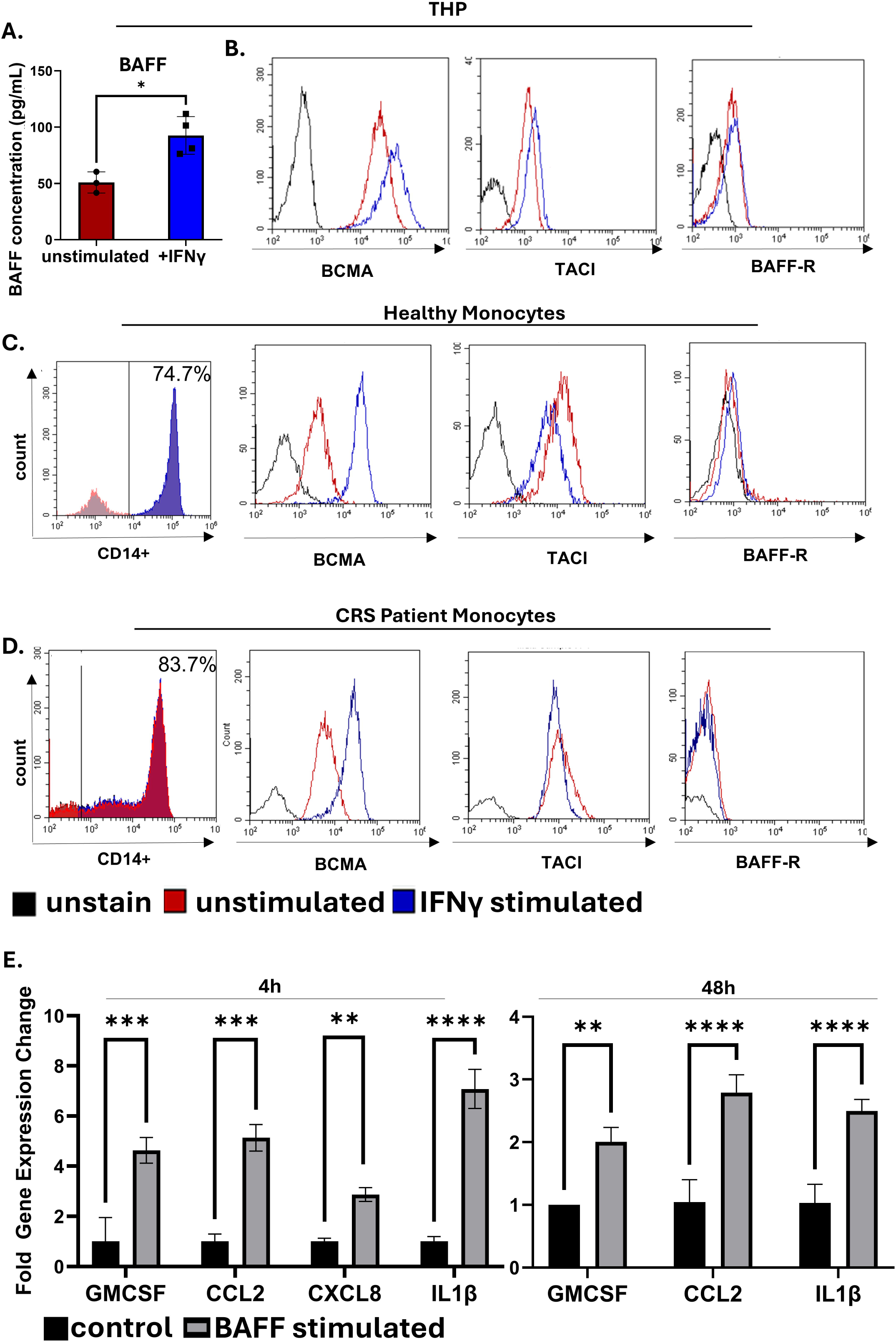
IFN-y stimulates BAFF secretion and upregulation of BCMA on monocytes. BAFF stimulates expression of CRS cytokines in monocytes. THP-1 were incubated with 50ng/ml rhlFNy. After 48h supernatant was collected and subject to (A) multiplex analysis to detect secreted BAFF (p=0.0122) (data represents 3 independent experiments) and (B) the cells were stained with a panel of antibodies for BCMA(PE), TACl(Pcy7), and BAFF-R (APC) and analyzed via flow cytometry. CD14+ monocytes were isolated from (C) healthy donor (n=6) or (D) CAR-T-CRS patient-derived (n=2) PBMCs and were stimulated and analyzed as in (A). E THP-1 cells were stimulated with 250ng/ml of rhBAFF. After various timepoints, RNA was isolated, and expression of various cytokines was measured by RT-qPCR (this data represents 2 independent experiments).

This was not surprising as monocytes are one of the main producers of BAFF, and that IFN-γ-induced BAFF has been shown in other contexts.^23,24^ We were then interested in whether monocytes express any receptors of BAFF, such that this secreted BAFF may act in an autocrine manner to stimulate monocytes to increase CRS cytokine production.

BAFF-R is known to be expressed at early stages of B-cell development, while TACI and BCMA are expressed on activated B-cells and plasma cells.^21^ In accordance with the pro-tumorigenic signals that BAFF produces, it is canonically known that various B-cell cancers have higher expression levels of these receptors than their healthy counterparts. The expression of these receptors is much less established on non-B-cell populations. We isolated the CD14+ monocyte populations from the peripheral blood mononuclear cells of healthy donors and CAR-T-treated patients who experienced CRS. We found that THP cells and monocytes from both healthy donors and CAR-T-CRS patients express BCMA and TACI. Furthermore, BCMA expression is substantially increased upon IFN-γ stimulation, potentially suggesting a mechanism of feedforward pro-inflammatory signaling (Figure 4B, C, D). The BAFF-receptor was not expressed on any monocyte population, and its expression was not affected by IFN-γ stimulation (Figure 4B, C, D).

### BAFF induces expression of CRS-related cytokines in monocytes

We then investigated whether BAFF could stimulate the expression of CRS-related cytokines in THP monocytes. We found that, at various timepoints, BAFF stimulation resulted in increased expression of cytokines in monocytes (Figure 4E). GM-CSF expression in monocytes increased two to fivefold at 48 hours and 4 hours, respectively. The expression of CCL2 increased at both timepoints, at three-to-five-fold times the unstimulated expression level. CXCL8 increased two-fold at 4 hours of BAFF stimulation. Lastly, IL-1β increased three to sevenfold over the baseline unstimulated expression level with BAFF stimulation at both timepoints.

### BAFF neutralization results in reduced cytokine secretion in vitro

Next, we investigated whether selective neutralization of BAFF with belimumab could decrease cytokine production in a CRS in vitro model. Belimumab is a recombinant monoclonal antibody directed against BAFF that is currently FDA-approved for the treatment of systemic lupus erythematosus.^25,26^ We utilized a tri-culture in vitro experimental setup that included CD19 CAR-T cells, monocytes (THP cells), and cancer cells (Raji or Jeko cells, representing Burkitt’s lymphoma and mantle cell lymphoma models). In the BAFF neutralization assay, cancer cells were pre-treated with belimumab for 24h before addition to co-culture, and cytokine release was measured after 16h of co-culture. We found that BAFF neutralization significantly diminished the production of multiple cytokines that are known to play a role in CAR-T-related CRS (Figure 5A). CCL2 was one CAR-T-CRS-related cytokine that was not significantly affected by BAFF neutralization.

**Figure 5.**
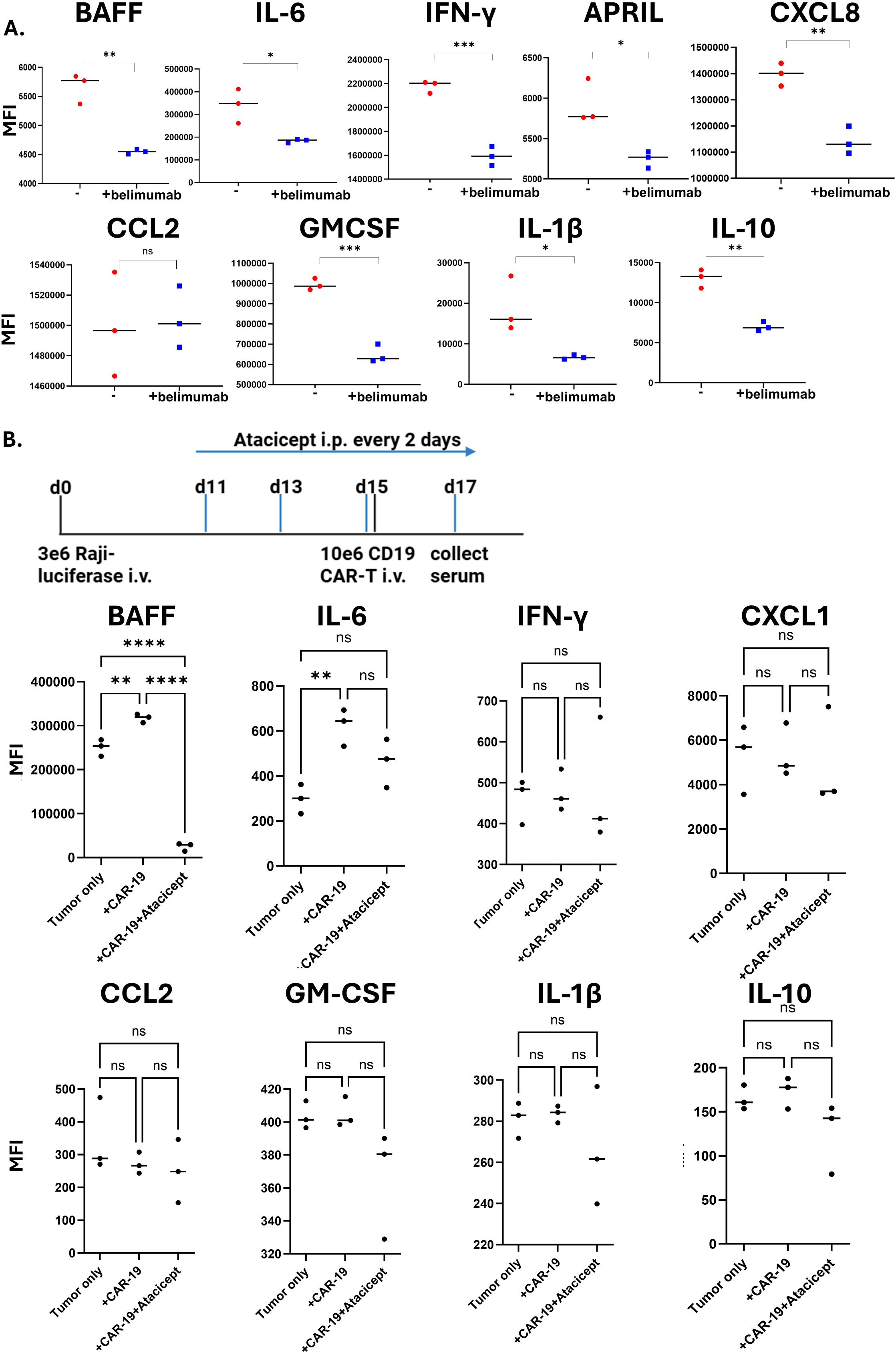
BAFF neutralization reduces cytokine release. (A) Jeko-1 cells were pre-incubated with 100µg/ml belimumab for 24h before co-culture with THP-1 and CD19 CAR-T cells (“-”: no belimumab). After another 16-24 the supernatant of the co-culture was harvested and subject to cytokine analysis by Multiplex. An unpaired t-test was used to compare untreated and belimumab conditions for each analyte. This data represents three independent experiments. P-values: BAFF 0.0017, IL-6 0.0234, IFN-γ 0.0004, APRIL 0.0161, CXCL8 0.0029, CCL2 0.88454, GM- CSF 0.0004, IL-1β 0.037, IL-10 0.0013. (B) SCIO beige mice were inoculated intravenously with 3 million Raji-luciferase cells. 3 mice were treated with atacicept every 2 days starting at day 11 post inoculation. At day 15 6 mice received 10 million CD19 CAR-T cells. Serum was harvested at day 17 and subjected to murine multiplex analysis.

### BAFF neutralization decreases CRS cytokine release in vivo

To investigate BAFF in CAR-T cell-related CRS in vivo, we inoculated scid-beige mice with 3 million Raji-luciferase cells intravenously. The experimental group received 100μg of atacicept every two days starting at day 11 post-inoculation. Atacicept is a fusion protein that consists of the binding portion of TACI and the Fc region of human IgG1 and neutralizes both BAFF and its related cytokine A proliferation-inducing ligand (APRIL).^27,28^ At day 15, 10 million CAR-T cells were injected intravenously into the +CAR-19 and +CAR-19+Atacicept groups (n=3 each). Murine cytokine levels were measured by multiplex analysis 2 days after CAR-T cell treatment. We found that BAFF was significantly elevated in the serum following CAR-T cell injection, supporting our hypothesis that BAFF plays a role in this pathophysiology. Further, cytokines that are critically involved in the CRS cascade, including IL-6, GM-CSF, and IL-10, trended downwards with BAFF blockade with atacicept (Figure 5B).

### BCMA neutralization results in reduced cytokine secretion in vitro

Because we found that BCMA is a receptor of BAFF that is both expressed at a baseline on monocytes and upregulated by IFN-γ, we were interested in whether BAFF may signal through BCMA to induce cytokine secretion in CRS. Similar to our BAFF neutralization assay with belimumab, THP cells were pre-incubated with a BCMA neutralizing antibody for 24h before addition to a co-culture with cancer cells and CD19 CAR-T cells. After 8-24 hours of co-culture, we observed a similar, significant reduction in CRS cytokines levels in vitro after pharmacological inhibition of BCMA, suggesting that BAFF may be signaling through BCMA on monocytes to produce cytokines involved in the CAR-T CRS mechanism (Figure 6).

**Figure 6.**
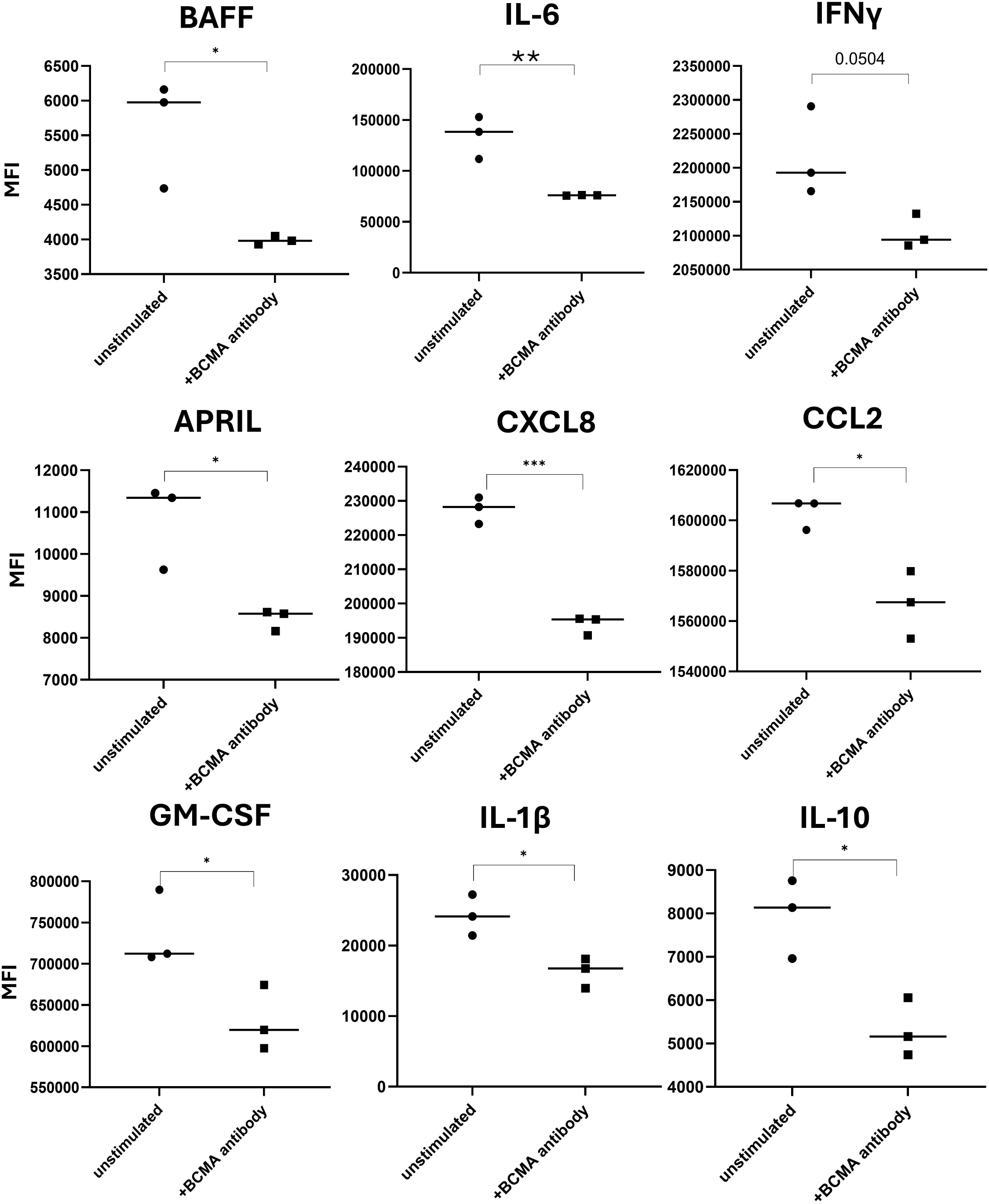
BCMA neutralization reduces cytokine release. THP-1 cells were pre-incubated with 10µg/ml of anti-BCMA neutralizing antibody for 24h. THPs were then co-cultured with Jeko-1 cells and CD19 CAR-T cells at a ratio of 5:1:5. Supernatant from the co-culture was harvested 8 or 24 hours and subject to cytokine analysis by Multiplex. This data represents 2 independent experiment. P-values: BAFF 0.0217, IL-6 0.0085, IFN-γ 0.0504, APRIL 0.0180, CXCL8 0.0002, CCL2 0.0128, GM-CSF 0.0388, IL-1β 0.0182, IL-10 0.0158.

### BAFF neutralization does not impair CAR-T cell function

Next, we investigated whether Belimumab as a potential treatment for CAR-T-related CRS would impair CAR-T cell function. First, we pre-incubated labeled Jeko-1 cells with belimumab for 24h before co-culture with CD19 CAR-T cells. After 16h of co-culture we stained all cells with propidium iodide (PI) and gated on cancer cells to measure CAR-T cell killing. Belimumab did not affect the cancer cell killing capability of CAR-T cells or have an appreciable impact on the health of either cell population alone (Figure 7A). CD19 CAR-T cells killed ∼93% of cancer cells with and without the addition of Belimumab to the co-culture. Belimumab only very slightly increased the PI+ CAR-T cells and cancer cells (11.6% to 13.2% of CAR-T cells and 24.8% to 26.6% of cancer cells without and with belimumab, respectively). When Atacicept was added to the co-culture of CD19 CAR-T cells and Raji cells, there was similarly no appreciable impact on CAR-T cell killing (59.2% and 56.5% killing without and with the addition of Atacicept, respectively) (Supplementary Figure 4A). Additionally, in the presence of monocytes, treatment with Belimumab did not impair the activation, as measured by CD3+CD69+ cell population, or degranulation, as measured by CD3+CD107a+ cell population after co-culture (Figure 7B and C respectively). 75.1% of CAR-T cells were activated without belimumab and 79.5% activated with the addition of belimumab in co-culture with monocytes and cancer cells (Figure 7B middle). Interestingly, the addition of belimumab to CAR-T cells alone significantly increased their activation level (15.4% without belimumab and 20.9% with the addition of belimumab) (Figure 7B, right). The addition of Atacicept had no effect on CAR-T cell activation in co-culture with Raji cells (77.2% without and 76.4% with Atacicept) (Supplementary Figure 4B). 42% of CAR-T cells were degranulated without Belimumab and 45.4% were degranulated with Belimumab in co-culture with cancer cells and monocytes (Figure 7C middle). Belimumab had no effect on the degranulation CAR-T cell alone without co-culture (19.0% without and 19.3% degranulated with belimumab) (Figure 7C right). Atacicept similarly had no effect on the degranulation status of CAR-T cells in co-culture with Raji cells (Supplementary Figure 4C). We conclude that BAFF neutralization as a strategy to curb CRS does not impact anti-tumor CAR-T cell ability.

**Figure 7.**
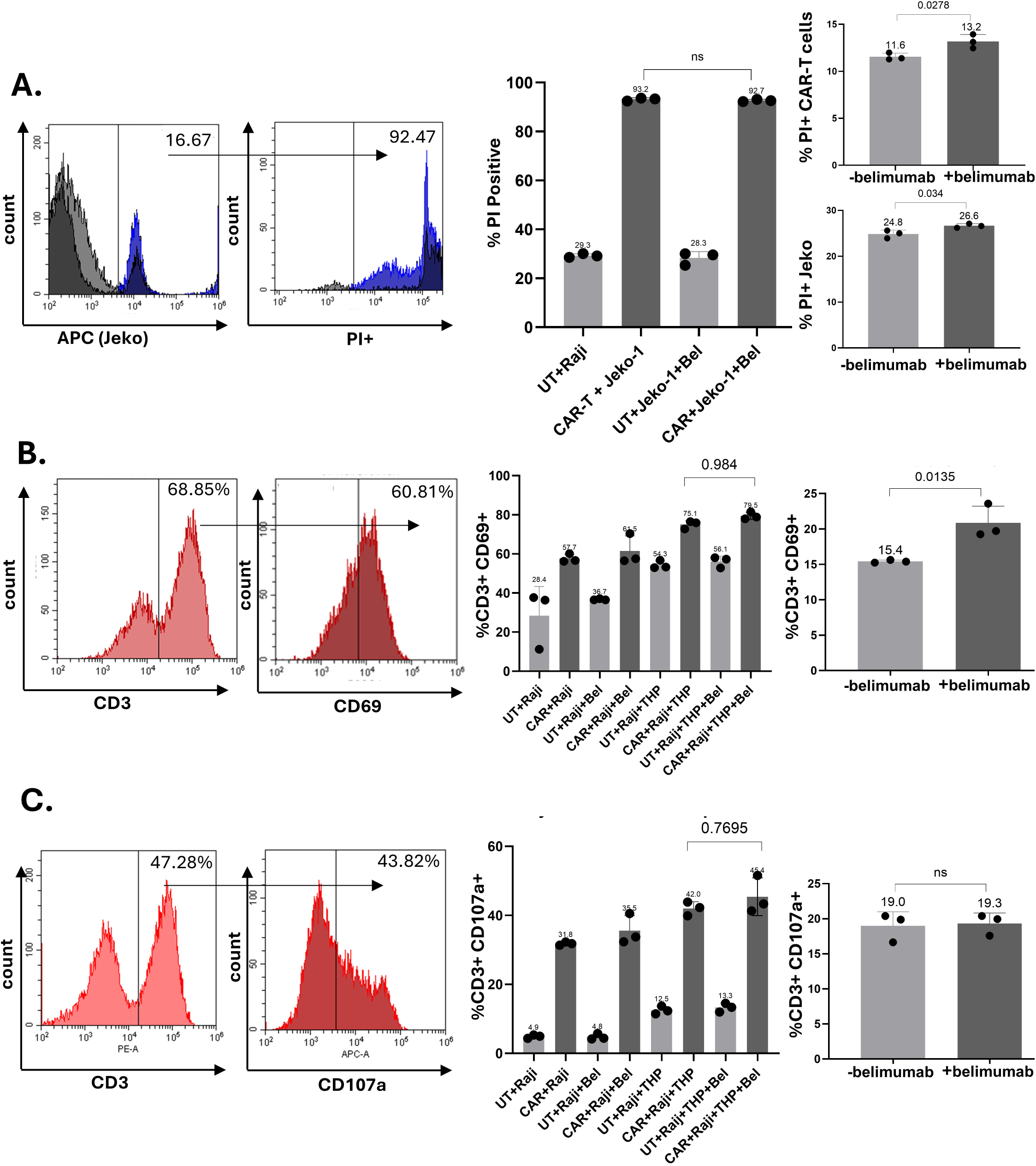
BAFF neutralization does not affect CAR-T cell function or activation. (A) Jeko-1 cells were stained with efluoro670 dye and pre-incubated with 100µg/ml belimumab (“Bel”) for 24h before co-culture with THP and CD19 CAR-T cells for an additional 16-20h. Cells were stained with propidium iodide and % APC and Pl - positive cells were reported. This data represents data from 2 CAR-T cell donors and 2 independent experimental replicates of each. (B) Raji cells were pre-incubated with 100µg/ml belimumab before co-culture with CAR-T and THP cells. After 16-20 additional hours of co-culture, cells were stained with CD3 (PE) and CD69 (Percp) and the percentage of CD3+ cells that are CD69+ was reported. (C) Raji cells were pre-incubated with 100µg/ml belimumab before co-culture with CAR-T and THP cells. Cells were immediately stained with CD107a (Pcy7). After 1 hour monensin golgi was added and cells were incubated for another 4-5 hours and were then stained with CD3 (PE). Percent positive CD3+ cells that stained with CD107a were reported for degranulation. The experiments in panels (B) and (C) represent 3 donors of CAR-T cells and 2 independent experiments per donor.

## Discussion

Challenges in CAR-T cell therapy involve side effects like CRS and ICANS, both of which result from cytokine release post CAR-T treatment.^18^ Though treatment with tocilizumab and corticosteroids is effective in many patients to treat side effects, it does not work in some patients.^29–31^ Further, tocilizumab is not effective for treating ICANS.^4,32^ Understanding the mechanism through which these cytokines are produced post-CAR-T therapy is crucial to developing better therapies to mitigate CRS and ICANS. This study aims to understand the unknown role of BAFF protein in mediating CRS and ICANS and how we can inhibit these toxicities related to CAR-T cell therapies. Our data show that BAFF is playing a major role in cytokine production by monocytes post-CAR-T activation, and that inhibiting BAFF might be a potential strategy to inhibit both CRS and ICANS.

We analyzed serum samples from patients who had CRS from 3 different CAR-T therapies and 3 different cancer diagnoses to ensure that our observations are not CAR-T or disease-specific. Patients who developed CRS showed an increase in BAFF levels in serum, while patients who did not experience CRS failed to show this elevation in BAFF. This observation intrigued us to explore further the role of BAFF in cytokine production post CAR-T therapy. Notably, in all the patients analyzed, serum BAFF levels were already elevated at baseline relative to the reported normal physiological concentration range. This was expected considering that B-cell cancer patients tend to have highly elevated BAFF levels in line with its pro-tumorigenic role.^20^ Since these patient samples are from three different CAR-T therapies, we are not able to compare tumor killing as there are significant differences among all the CAR-T products including dosing, expansion and killing kinetics. We are considering only CRS as a criteria in these analysis.

Our data also shows an increase in serum APRIL, a cytokine that is analogous to BAFF and shares the BCMA and TACI receptors, in CAR-T-CRS patients.^33^ We observed expression of both these receptors, which are generally considered B-cell lineage specific, on monocytes. We were compelled to focus on BAFF because APRIL levels are very low compared to BAFF and inhibition of BAFF specifically using Belimumab inhibited the secretion of cytokines. Monocytes are known to play a key role in CRS; hence, expression of BAFF and its receptors on monocytes warrants a detailed study on autocrine BAFF signaling in these cells and its functional implications. Further, BCMA expression on monocytes was upregulated after exposure to IFN-γ. We focused on IFN-γ in this study because of its prominent contribution to myeloid activation and its established role in CAR-T cell-related CRS.^34,35^ BCMA neutralizing antibody treatment also resulted in significant inhibition of cytokines in a tri-culture assay of tumor cells, CAR-T cells, and monocytes, which implies that the upregulated BCMA expression after exposure to IFN-γ might be a key upstream event for cytokine production related to CRS and ICANS. BAFF-induced NF-ĸB signaling pathways might be contributing to the production of these cytokines, as inhibiting BCMA or BAFF results in decreased levels of cytokines. NF-ĸB is a transcription factor that regulates the expression of most of these pro-inflammatory cytokines.^36–39^ BAFF signaling through BCMA is known to activate classical NF-kB signaling, which might be inducing the secretion of these pro-inflammatory cytokines.^40–42^

In support of other reports, our data show that monocytes are the key cytokine producers and cellular mediators of CAR-T-CRS.^3,4,9,35^ BAFF is produced by monocytes in the tri-culture system of CAR-T cells, tumor cells, and monocytes. The increase in BAFF observed in these cancer patients post-CAR-T therapy might be contributed to by monocytes, as evident from our data. Inhibiting BAFF resulted in a significant reduction of cytokines involved in both CRS (IL-6, CCL2, CXCL8, IFN-γ) and ICANS (IL-1β, GM-CSF, and IL-15) in the in vitro tri-culture system. Although BAFF has not previously been investigated in the context of CAR-T-related CRS, our findings are supported by previous work demonstrating BAFF-induced expression or secretion of multiple relevant cytokines, including IL-6, IL-1β, and IL-10 in different disease models.^43–45^

BAFF autocrine signaling resulting from BAFF binding to BCMA or TACI on monocytes might be a contributing factor to these spiked cytokine levels post-CAR-T therapy, responsible for CRS and ICANS. Our data propose that inhibiting BAFF ahead of CAR-T therapy will be a potential way to inhibit cytokine production and associated toxicities. As Belimumab is a specific inhibitor for human BAFF, we used Atacicept, which is capable of inhibiting both murine BAFF and APRIL, in mouse experiments. Though mouse models to mimic CAR-T-related CRS are reported, an unrealistically high number of CAR-T cells per kilogram body weight is required to induce CRS in these mice.^3^ Hence, murine models are not the optimal models to mimic human CRS, but our in vivo data show that Atacicept treatment resulted in inhibition of many murine cytokines after CAR-T therapy. A recent study reported that GM-CSF neutralization inhibits CRS and neurotoxicity after CD19 CAR-T cell therapy.^8^ Interestingly, we observed GMCSF inhibition after treatment with Belimumab, which points to the possibility that BAFF might be upstream of these cytokines, including GMCSF.

When identifying the contribution of the three cell types to the release of CRS cytokines, we found that the interaction of cancer cells with CAR T-cells most preferentially induced GM-CSF, IFN-γ, and IL-10 secretion. This is consistent with the working model of CAR-T-related CRS, in which T-cells activated by tumor antigen binding secrete high levels of GM-CSF and IFN-γ, which then stimulate additional cytokine secretion from monocytes.^4^ The combination of CAR-T and THP cells resulted in the greatest increase of IL-6, CCL2, CXCL8, and IL-1β out of the other cell combinations, also supporting the current understanding that activated CAR-T cells stimulate monocytes to secrete these cytokines. Lastly, we found that monocytes produce higher levels of BAFF than cancer cells or CAR-T cells at baseline, and the combination of CAR-T and THP cells yielded the greatest increase in BAFF levels out of these combinations. Overall, these results are in accordance with the pathophysiology proposed in the literature and our hypothesis that cytokines derived from activated CAR-T cells can induce monocytes to secrete various pro-inflammatory cytokines, including BAFF.^4,17^

Importantly, Belimumab or Atacicept treatment did not affect CAR-T cytotoxic function or viability. Belimumab is already FDA-approved for systemic lupus erythematosus, so it might be a viable therapeutic approach to add Belimumab to current CRS/ICANS treatment strategies as a combination therapy or as a monotherapy to better mitigate CRS in patients.^25^ We anticipate inhibition of ICANS in Belimumab-treated patients as our in vitro and in vivo data point to decreased levels of cytokines involved in ICANS, such as IL1β, IL10, GMCSF, IL-6, and IL-15^9,46–48^. Our data strongly suggest that inhibiting BAFF will potentially modulate cytokine secretion by monocytes and will help to control complications like CRS and ICANS post-CAR-T cell therapies.

## Methods

### Patient sample collection

Serum and plasma samples were obtained from CAR-T-treated patients who received commercial Axicabtagene Ciloleucel at University Hospitals in Cleveland OH, LMY-920 as part of a clinical trial at University Hospitals, or KURE-19 as part of a clinical trial.

### Cell lines

Raji, Jeko-1, and THP-1 cells were purchased from the American Type Culture Collection (ATCC). Both cell lines were cultured in HyClone™ RPMI 1640 Media (Cytiva, catalog #: SH3025501) supplemented with 10% fetal bovine serum (Sigma Aldrich, catalog # F0926) and 1% penicillin streptomycin and incubated in 5% CO_2_ at 37°C. For in vivo experiments, Raji or Jeko-1 cells were modified by lentiviral transduction to stably express the firefly luciferase gene using a pLM expression vector (Raji-luc).

### Primary cells and CAR-T cells

Human peripheral blood mononuclear (hPBMCs) cells were isolated from de-identified healthy volunteers from the CWRU Hematopoietic Biorepository and Cellular Therapy Core under a CWRU IRB-approved protocol (IRB#09-90-195). Donor blood was diluted 1:1 with 1X PBS and layered over Ficoll-Paque for density gradient separation. The samples were centrifuged at 400xg for 35 minutes. Plasma was removed, and the PBMC monolayer was isolated. Primary monocytes were isolated using the Mojosort^TM^ Human CD14+ Monocyte Isolation Kit following the manufacturer’s protocol (Biolegend, catalog # 480047). Monocyte purity was tested by staining for CD14 (Pcy7 Biolegend, catalog).

CD19 CAR-T cells that were used for both in vitro and in vivo experiments were sourced from the CWRU Hematopoietic Biorepository and Cellular Therapy core. The CAR-T cells were harvested by T-cell apheresis 8 days after intravenous administration.

### Luminex assay

Luminex is a multiplex platform that analyzes multiple cytokines at the same time, using a validated platform. Each analyte has a bead that contains a fluorophore. The beads are run together and followed with streptavidine-phycoerthrin (SA-PE) detection antibodies. The dual laser system detects the bead fluorescence identification value, which is selected to be at least 50 beads, and the secondary SA-PE detection, quantified against a standard curve analyzed using the same complexity in the assay. Samples were obtained from patients and aliquoted according to manufacturing guidelines. Each sample was run in duplicate for each analysis of 50 beads, as is the standard curve. Quantitative values are analyzed based on the mean fluorescent intensity of the bead and SA-PE, and the pg/ml concentration of the standard curve.

### Cytokine multiplex assay

LEGENDplex^TM^ multiplex, bead-based flow cytometry assays were used to analyze in vitro and in vivo cytokine release (BioLegend). The human custom kit included IL-6, IL-8, IL-10, MCP-1, IL-1β, IFN-γ, APRIL, BAFF, GM-CSF, and the murine analytes included IL-6, IL-8, IL-10, MCP-1, IL-1β, IFN-γ, BAFF, and GM-CSF. For in vitro cytokine analyses, cell cultures were centrifuged, and the supernatant was stored at -80°C. The manufacturer’s protocol was followed for sample preparation and analyzed using the CytoFLEX (Beckman Coulter). For serum analyses, mice were bled by tail vein into EDTA-coated tubes (Sarstedt, catalog # NC9954576) and centrifuged at 2600 x rpm for 15 minutes. Serum was aliquoted to avoid freeze-thaw cycles and stored at -80°C until used.

### Analysis of cell types contributing to cytokine release

Jeko-1 cells, CD19 CAR-T cells, and THP-1 cells were cultured alone or in every combination at a 1:1:1 ratio (100,000 cells each) for 24h, when the supernatant was harvested for cytokine analysis by Multiplex as described in Cytokine Multiplex Assay.

### Monocyte receptor staining

To assess BAFF receptor expression on primary monocytes, 1μL of each antibody (BAFF-R:APC, TACI:Pcy-7, BCMA:Percp, Biolegend catalog #’s: 316916, 311908, 357510) was added to 50μL of cells suspended in PBS and incubated at room temperature for 15 minutes in the dark. 1mL of PBS was added, cells were centrifuged, and unbound antibody was removed. The stained monocytes were resuspended in 150μL of PBS and run on the CytoFLEX flow cytometer (Beckman Coulter).

### IFN-**γ**-induced BAFF and receptors experiment

To assess IFN-γ stimulation of BAFF from monocytes and receptor expression, THP cells were stimulated with 50ng/mL of recombinant human IFN-γ (Peprotech, catalog #: 300-02) for 48 hours. The cells were processed for flow cytometry to detect IFN-γ-stimulation of receptor expression (see Monocyte receptor staining). The supernatant was saved for BAFF cytokine analysis using a multiplex assay (see Cytokine multiplex assay).

### Real-time quantitative PCR (RT-qPCR)

RNA was harvested from BAFF-stimulated hPBMCs or THP cells using the RNAEasy RNA extraction kit (Qiagen, catalog #: 74014) following the manufacturer’s protocol.

RNA purity and concentration were measured using the NanoDrop™ 2000/2000c spectrophotometer (Thermo Scientific). Only samples with an A260/A280 ratio of between 2.0 to 2.1 were utilized for reverse transcription and qPCR. 1μg of RNA of each condition was combined with Script^TM^ Reverse Transcription Supermix (Bio-Rad, catalog# 1708840) and nuclease-free water. The RNA was reverse transcribed to cDNA using the protocol: 5 min at 25 °C, 20 min at 46 °C, and 1 min at 95 °C in a T100 Thermocycler (Bio-Rad, catalog #: 1861096). For qPCR, cDNA was combined with iQ Sybr green Supermix (Bio-Rad, 170880), nuclease-free water, and forward/reverse primers (see Table.1 for sequences) in a 96-well plate. The plate was loaded into a C1000 Touch^TM^ Thermal Cycler (Bio-Rad) and the manufacturer’s instructions were followed for data acquisition and analysis.

**Table 1:**
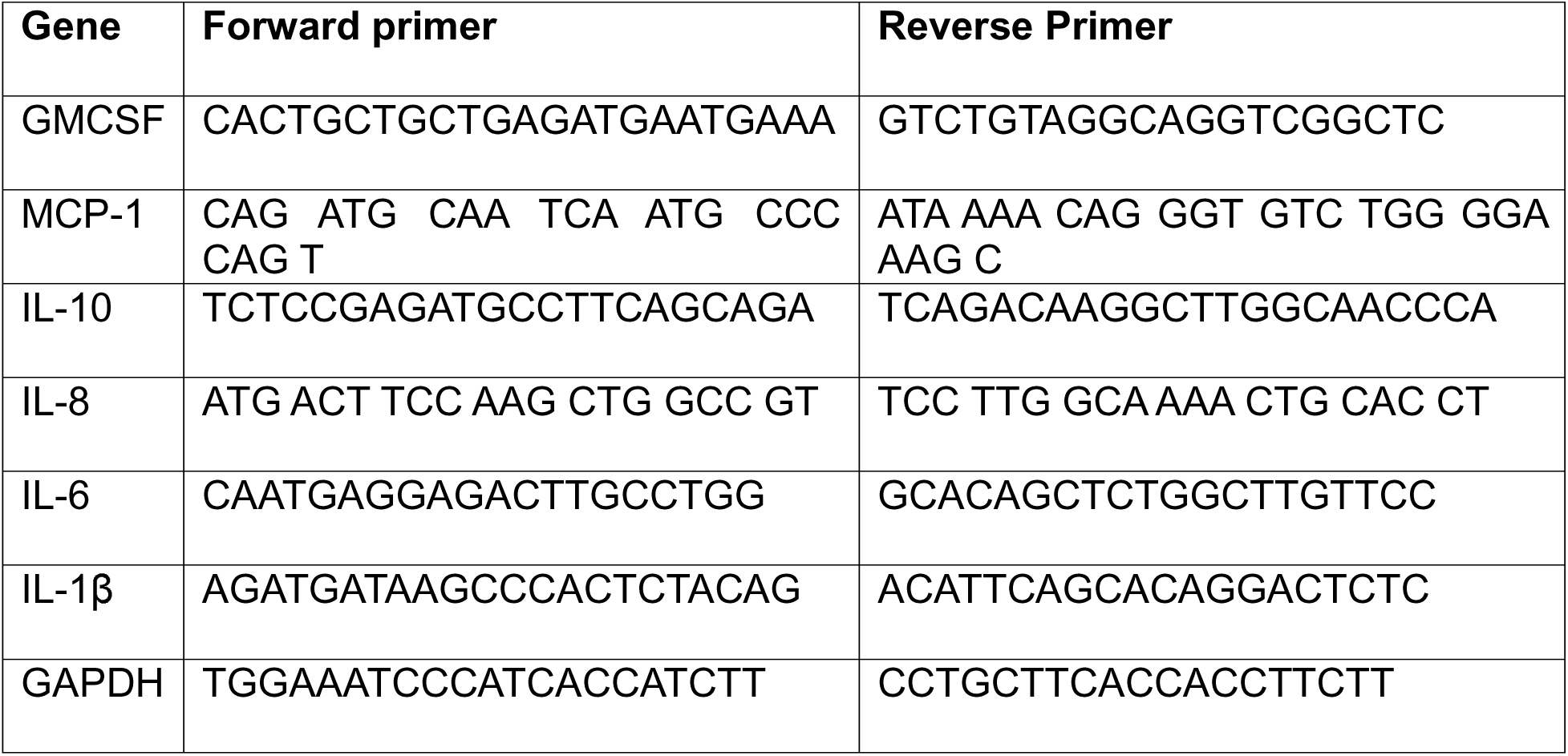
Primer sequences used in RT-qPCR.

### BAFF & BCMA neutralization experiments

Belimumab is a humanized IgG1λ monoclonal antibody against B-cell activating factor that antagonizes BAFF activity. For these experiments, Belimumab was purchased from MedchemExpress (catalog#: HY-P9952) and used at a dose of 100μg/ml in vitro. Jeko-1 cells were pre-incubated with Belimumab for 24h before co-culture with THP-1 and CD19 CAR-T cells (5:1:5 ratio CAR-T: Cancer: THP cells). After another 16-24h the supernatant of the co-culture was harvested and subject to cytokine analysis by Multiplex, as described in Cytokine Multiplex Assay. The same procedure was followed for the BCMA neutralization experiments, but 10μg/ml of the anti-BCMA neutralizing antibody (Acro Biosystems, catalog #: BCA-M43) was utilized instead of Belimumab.

### In vivo modeling of CRS

All in vivo experiments were approved by the CWRU IACUC and were conducted in accordance with IACUC protocol number 2016–0307. The mice were housed in the Small Animal Imaging Facility (SAIF) in University Hospitals in a pathogen-free environment. The SAIF is equipped with controlled environmental controls, including proper temperature, humidity, and lighting, as well as HEPA-filtered air circulation. Mice were inspected daily by staff technicians, and cages and bedding were changed on a weekly basis. To model CAR-T-related CRS in vivo, SCID-Beige mice (Taconic Biosciences, C.B-*Igh*-1b/GbmsTac-Prkdc*^scid^*-Lyst^bg^N7) were inoculated intravenously with 3 million Raji-luciferase cells. 11 days after inoculation 3 mice received treatment with Atacicept. The experimental group received 100μg of atacicept intraperitoneally every other day through the remainder of the experiment. At day 15, 6 mice (n=3 with and without Atacicept treatment) received 10 million CAR-T cells intravenously. At day 17, serum was collected and subject to murine multiplex cytokine analysis.

### T-cell functional assays

#### Cytotoxicity assay

Jeko-1 or Raji cells were labeled with eFlour670^TM^ Cell Proliferation Dye (ThermoFischer Scientific, catalog #: 65-0840-85) and pre-incubated with 100μg/mL belimumab for 24h. CD19 CAR-T cells were then co-cultured with THP monocytes and labeled/pre-treated Jeko-1 cells at a ratio of 5:1:5 (100,000 CAR-T:20,000 THP:100,000 Raji) in complete RPMI-1640 media containing 2ng/mL of IL-2 and additional belimumab. After 16h of co-culture, the cells were stained with propidium iodide (PI, Cell Signaling Technology, 4087S), and cancer cell death was measured by the APC (eFlour670) + PE (PI)+ cell population. The supernatant of the co-culture experiments was saved for cytokine analysis by multiplex.

#### Activation assay

Raji cells were pre-treated with 100μg/mL belimumab for 24h and cultured with CD19 CAR-T cells and THP monocytes at a ratio of 5:1:5 (150,000 CAR-T: 30,000 Raji cells: 150,000 THP) in complete RPMI-1640 media containing 2ng/mL of IL-2. After 16h of co-incubation, the cells were stained with PE:CD3 (Biolegend, 300408) and Percp: CD69 (Biolegend, catalog#: 310926). T-cells were gated on by CD3 positivity and the percentage of CD69 expressing cells on flow cytometry was reported.

#### Degranulation assay

Raji cells were pre-treated with 100μg/mL belimumab for 24h and cultured with CD19 CAR-T cells and THP monocytes at a ratio of 5:1:5 (150,000 CAR-T: 30,000 Raji cells: 150,000 THP) in complete RPMI-1640 media containing 2ng/mL of IL-2. The cells were immediately stained with Pcy7 or APC CD107a (Biolegend, catalog#: 328618, 328619).

After 1 hour, GolgiStop protein transport inhibitor (BD Biosciences) was added. Following another 5 hours of incubation, the cells were labeled with PE: CD3 antibody, and degranulated T-cells were measured by the percentage of CD3+ T-cells that expressed CD107a.

### Statistical analyses

Unpaired students t-tests were used to compare cytokine levels in untreated and treated groups (unstimulated versus IFN-γ-stimulated THP cells in Figure 4, untreated versus Belimumab or BCMA-neutralizing antibody treatment in Figures 5,6). A one-way ANOVA with Tukey’s multiple comparison correction was utilized to compare the means of the T-cell functional assays and in vivo cytokine levels (Figures 5, 7 and Supplementary Figure 4).

## Supporting information

Supplementary Combined

## Acknowledgements

We would like to acknowledge and thank Tracey Bonefield and David Fletcher of the CWRU Bioanalyte Core for their assistance with the planning and data acquisition of cytokine levels in CAR-T cell patients. We would further like to acknowledge Jane Reese and Kelao Neumbo of the CWRU Hematopoetic Biorepository and Cellular Therapy Core for kindly providing CD19 CAR-T cells for use in vitro and in vivo experiments. Additionally, Luminary Therapeutics is sponsoring the clinical trial of LMY-920-001 (NCT05312801) and provided serum from a CAR-T treated patient who experienced CRS that we included in our serum cytokine analysis. Biorender.com was used to create the graphical abstract.

## Author contributions

C.F. performed the experiments, analyzed data, and wrote the manuscript, D.W. is the sponsor and IND holder of clinical trial, LM and PC are Principal Investigators of clinical trials from whom we obtained patient samples for this study. R.P. conceived the study, oversaw the study, designed experiments, guided research personnel, performed data analysis, and edited the manuscript.

## Data availability statement

The authors confirm that the data supporting the findings of this study are available within the article.

