## Supplementary figures and images for "B-cell activating factor plays a critical role in CAR-T cell-associated cytokine release syndrome"

### Supplementary Figure-1

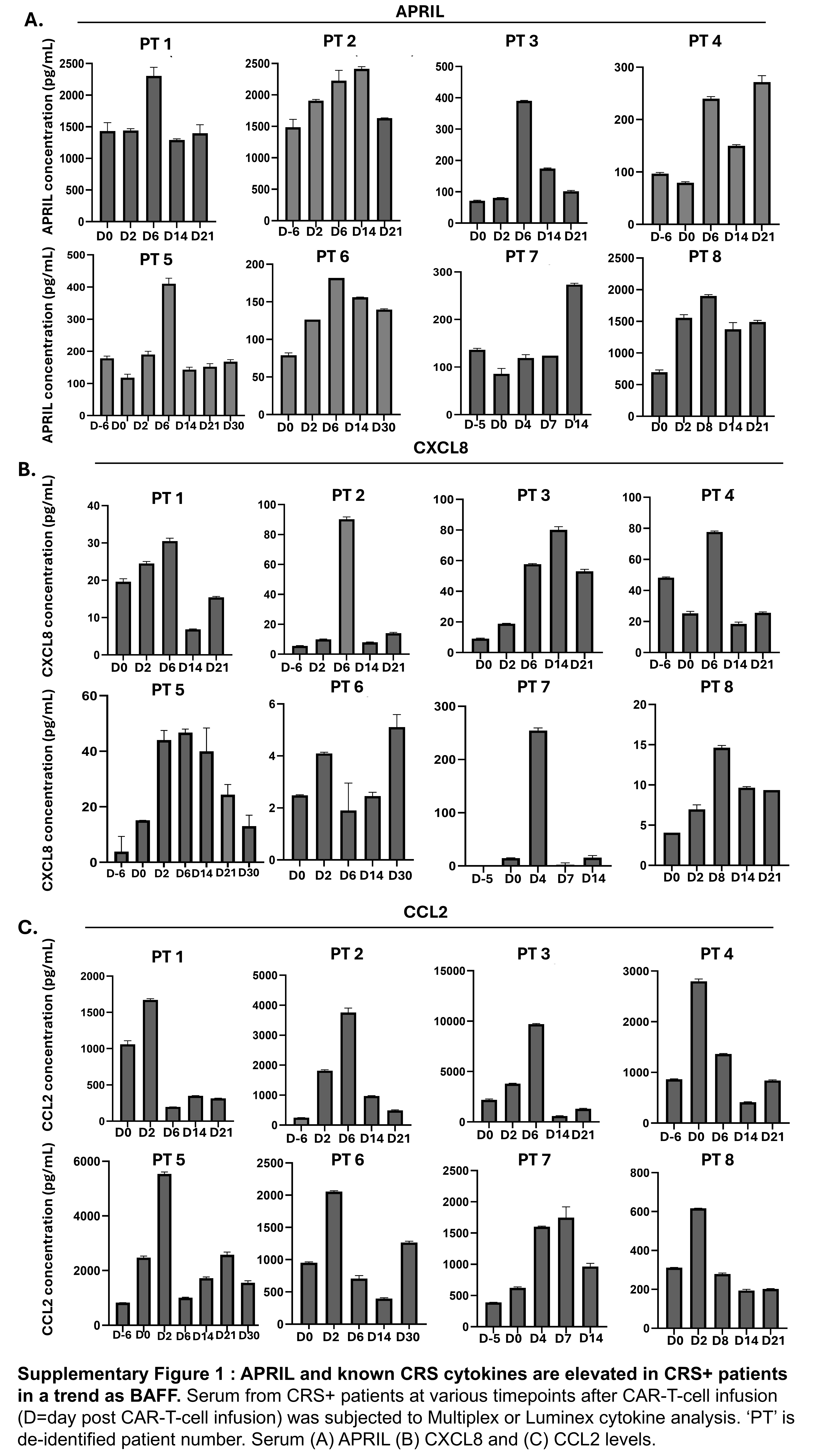

### Supplementary Figure-2

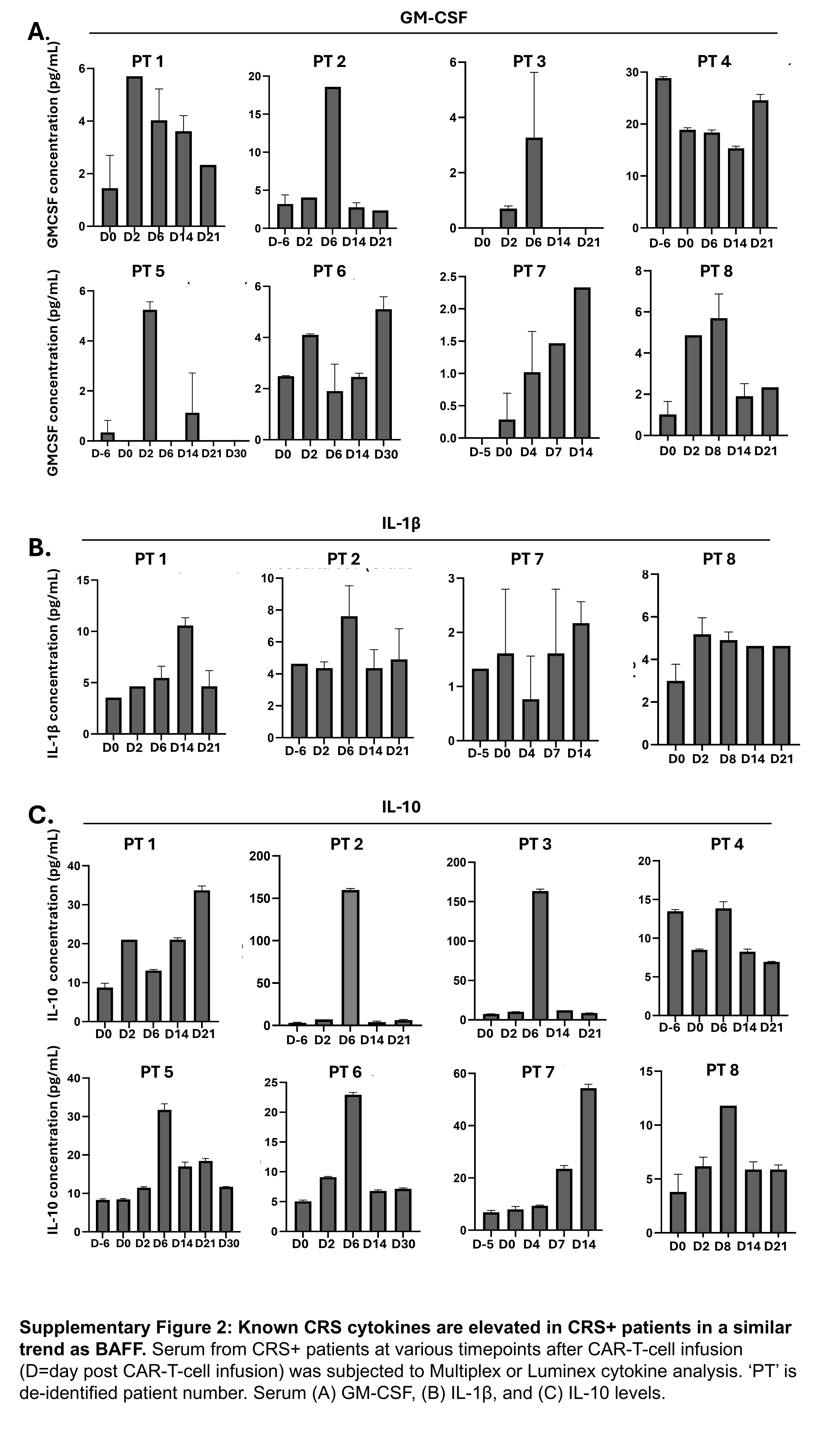

### Supplementary Figure-3

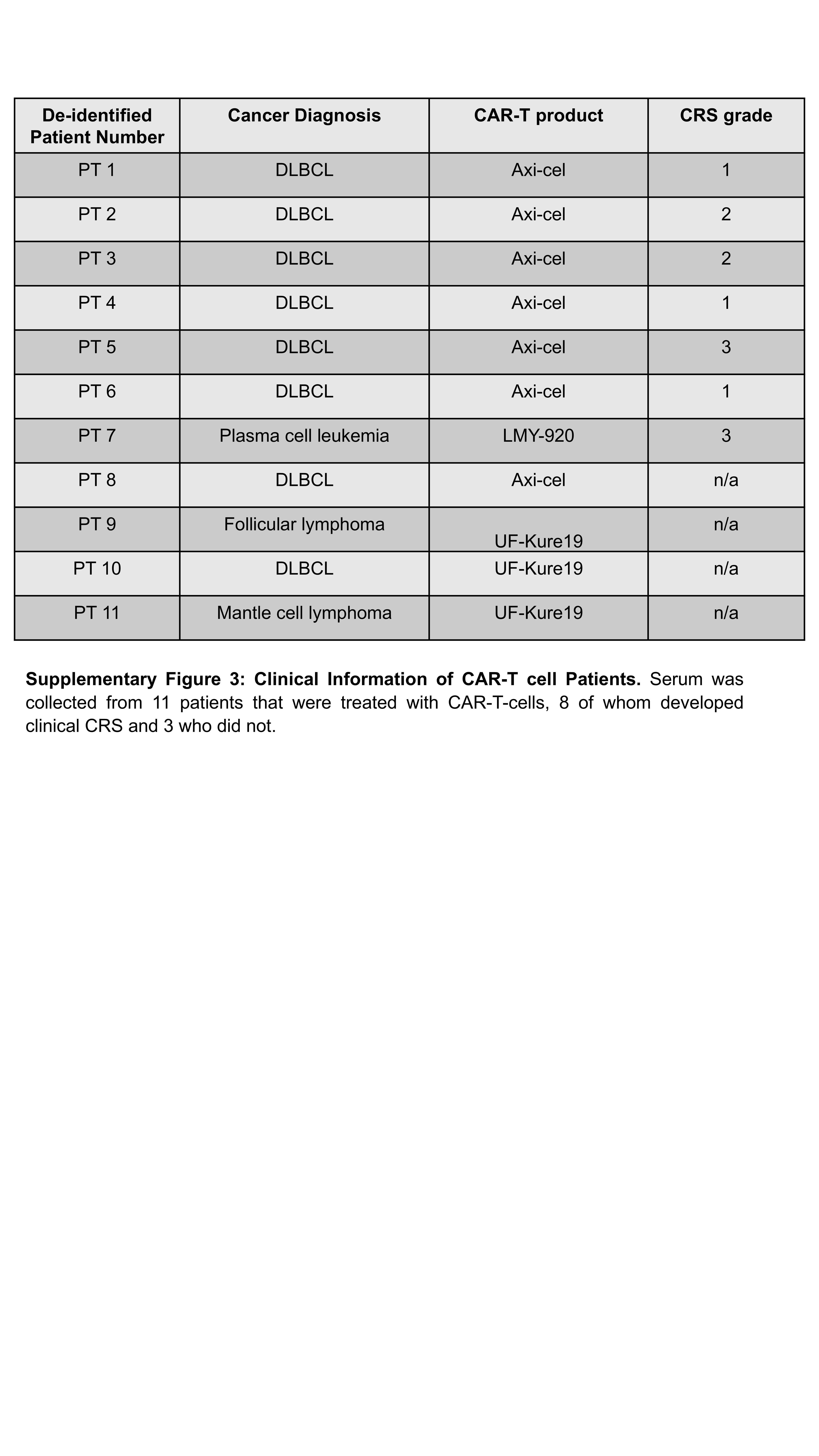

### Supplementary Figure-4

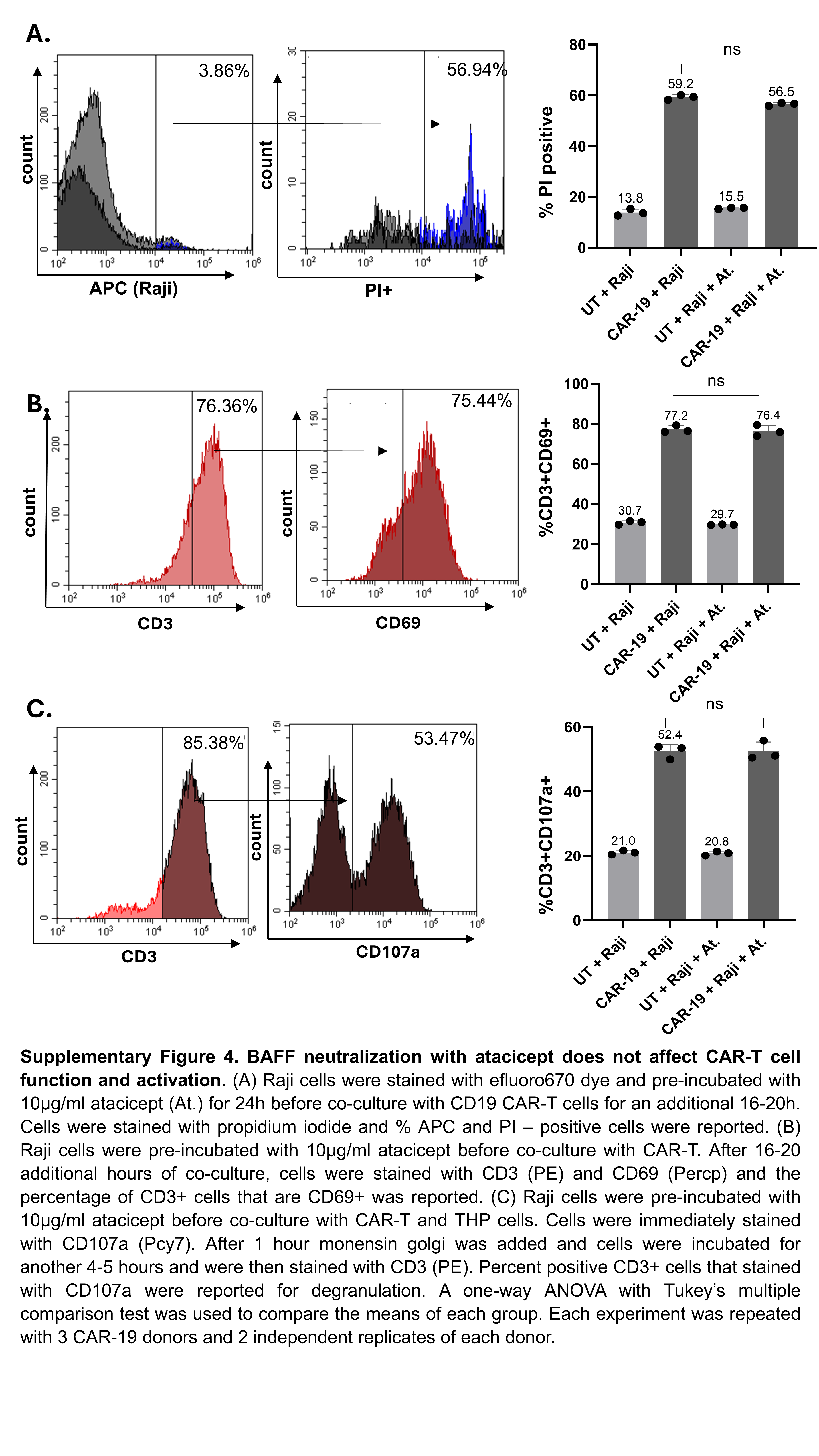
