## Supplementary Combined for "B-cell activating factor plays a critical role in CAR-T cell-associated cytokine release syndrome"

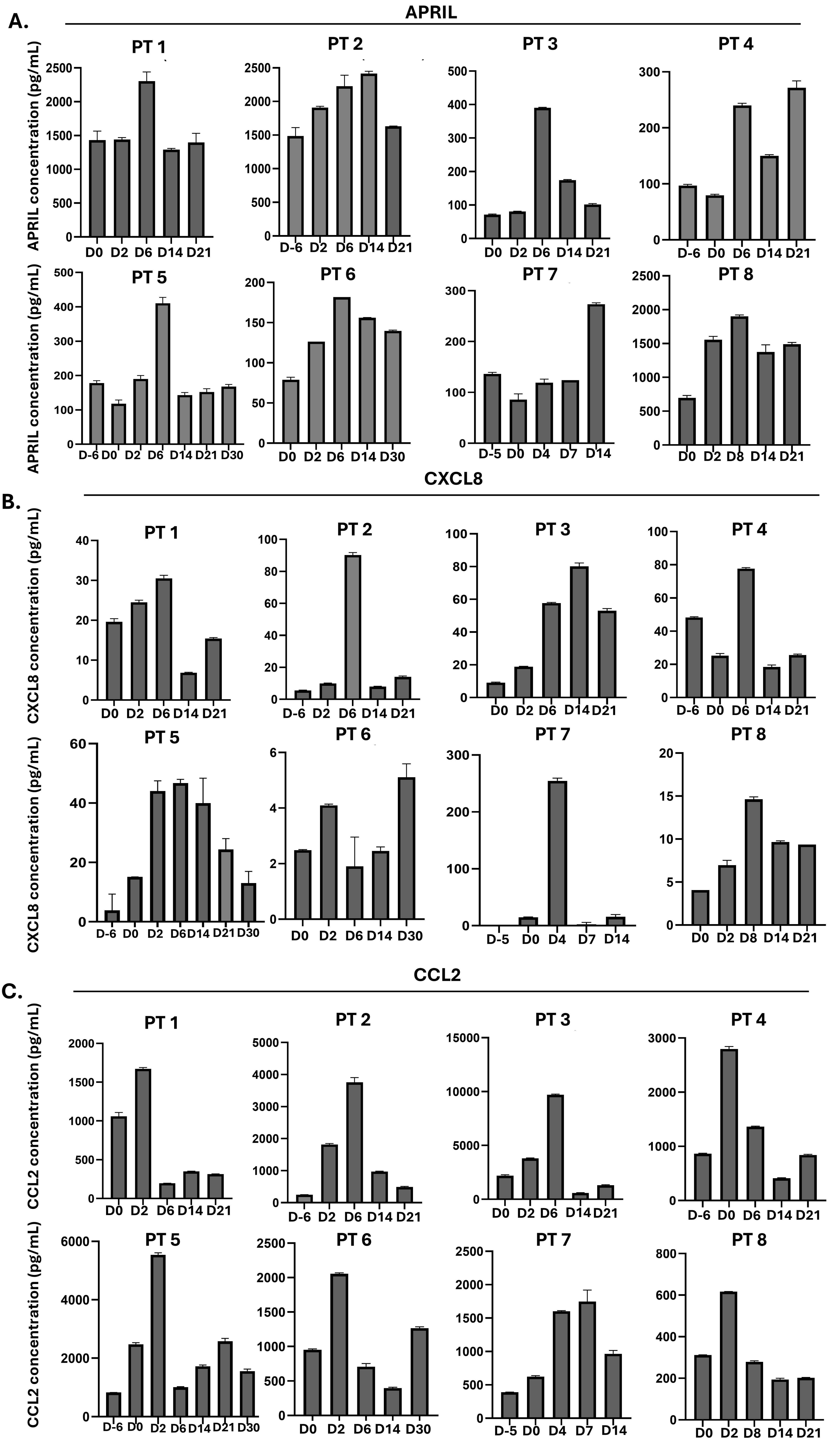


**Supplementary Figure 1 : APRIL and known CRS cytokines are elevated in CRS+ patients in a trend as BAFF.** Serum from CRS+ patients at various timepoints after CAR-T-cell infusion (D=day post CAR-T-cell infusion) was subjected to Multiplex or Luminex cytokine analysis. 'PT' is de-identified patient number. Serum (A) APRIL (B) CXCL8 and (C) CCL2 levels.

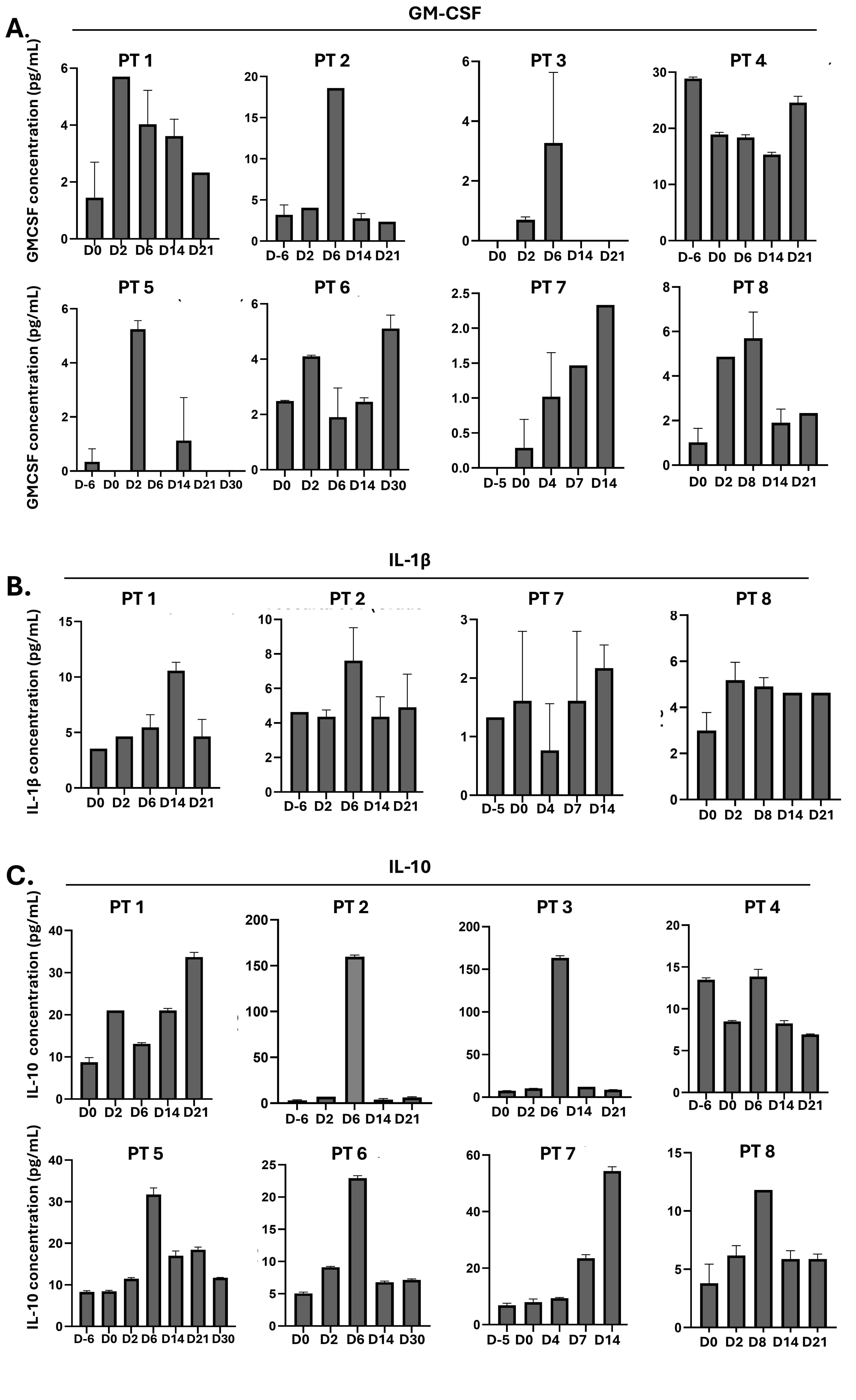


**Supplementary Figure 2: Known CRS cytokines are elevated in CRS+ patients in a similar trend as BAFF.** Serum from CRS+ patients at various timepoints after CAR-T-cell infusion (D=day post CAR-T-cell infusion) was subjected to Multiplex or Luminex cytokine analysis. ‘PT’ is de-identified patient number. Serum (A) GM-CSF, (B) IL-1β, and (C) IL-10 levels.

| De-identified Patient Number | Cancer Diagnosis | CAR-T product | CRS grade |
|---|---|---|---|
| PT 1 | DLBCL | Axi-cel | 1 |
| PT 2 | DLBCL | Axi-cel | 2 |
| PT 3 | DLBCL | Axi-cel | 2 |
| PT 4 | DLBCL | Axi-cel | 1 |
| PT 5 | DLBCL | Axi-cel | 3 |
| PT 6 | DLBCL | Axi-cel | 1 |
| PT 7 | Plasma cell leukemia | LMY-920 | 3 |
| PT 8 | DLBCL | Axi-cel | n/a |
| PT 9 | Follicular lymphoma | UF-Kure19 | n/a |
| PT 10 | DLBCL | UF-Kure19 | n/a |
| PT 11 | Mantle cell lymphoma | UF-Kure19 | n/a |

**Supplementary Figure 3: Clinical Information of CAR-T cell Patients.** Serum was collected from 11 patients that were treated with CAR-T-cells, 8 of whom developed clinical CRS and 3 who did not.

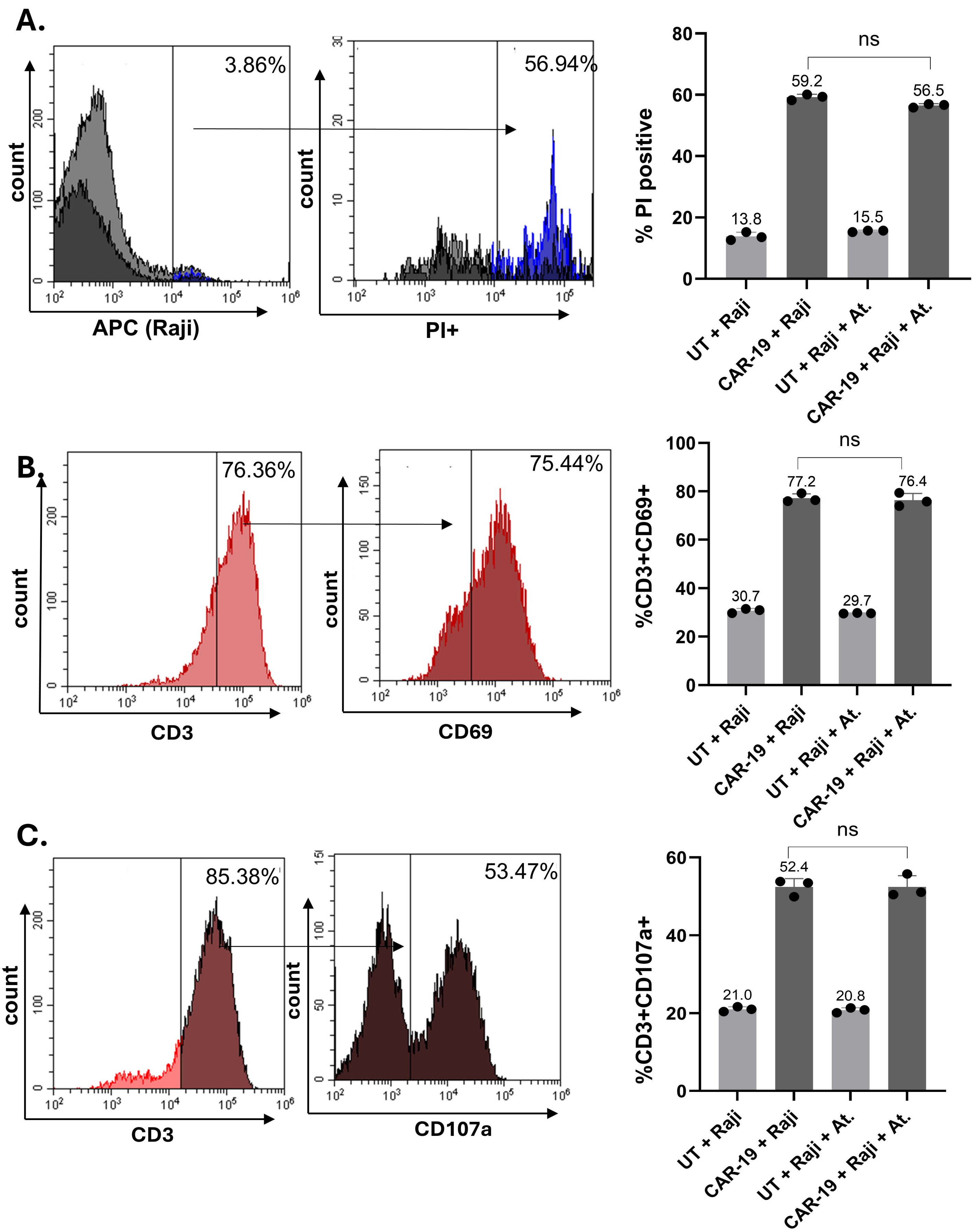


**Supplementary Figure 4. BAFF neutralization with atacicept does not affect CAR-T cell function and activation.** (A) Raji cells were stained with efluoro670 dye and pre-incubated with 10μg/ml atacicept (At.) for 24h before co-culture with CD19 CAR-T cells for an additional 16-20h. Cells were stained with propidium iodide and % APC and PI – positive cells were reported. (B) Raji cells were pre-incubated with 10μg/ml atacicept before co-culture with CAR-T. After 16-20 additional hours of co-culture, cells were stained with CD3 (PE) and CD69 (Percp) and the percentage of CD3+ cells that are CD69+ was reported. (C) Raji cells were pre-incubated with 10μg/ml atacicept before co-culture with CAR-T and THP cells. Cells were immediately stained with CD107a (Pcy7). After 1 hour monensin golgi was added and cells were incubated for another 4-5 hours and were then stained with CD3 (PE). Percent positive CD3+ cells that stained with CD107a were reported for degranulation. A one-way ANOVA with Tukey's multiple comparison test was used to compare the means of each group. Each experiment was repeated with 3 CAR-19 donors and 2 independent replicates of each donor.
